# Pan-cancer HLA Gene-mediated Tumor Immunogenicity and Immune Evasion

**DOI:** 10.1101/2021.05.17.444511

**Authors:** Xutong Gong, Rachel Karchin

## Abstract

Human Leukocyte Antigen (HLA) expression contributes to the activation of anti-tumor immunity through interactions with T cell receptors. However, pan-cancer HLA-mediated immunogenicity and immunoediting mechanisms has not been systematically studied. In a retrospective analysis of 33 tumor types from the Cancer Genome Atlas, we uncovered HLA class I and class II differential expression, which outperformed traditional clinical metrics in predicting patient survival. We also characterized the distribution of HLA supertypes across cancers and showed that patients with high HLA allelic diversity and gene expression had better prognosis. Immune microenvironments with varied survival outcomes could be predicted using a neural network model trained on HLA expression data. Furthermore, we identified a subset of tumors which upregulated HLA class I but not class II genes and exploited HLA-mediated escape strategies. Our results suggest the potential of using HLA expression data to predict patient prognosis. Taken together, we emphasize the crucial role of HLA upregulation in shaping prolonged anti-tumor immunity.

**Synopsis:** In a retrospective analysis of 11080 patients of 33 TCGA cancer types, we showed HLA class I and class II differential expression shape various immune microenvironments and tumor immunoediting mechanisms, predicting tumor immunogenicity and patient survival.

## Introduction

The tumor microenvironment (TME) is a hostile environment for infiltrating immune cells and poses significant challenges to their proper function. For instance, hypoxia and nutritional depletion hinder lymphocyte viability and dampen the immune response against tumors (1). Tumors may exploit immunosuppressive strategies to evade immune rejection, including the expression of immune checkpoints like programmed death-ligand 1 (PD-L1) and the recruitment of immune suppressor cells (2,3). Shaped by the complex interactions between cellular components in the TME, the immune microenvironment varies significantly among patients and across tumor types.

In recent years, significant efforts have been made to study the dynamic components in the TME to shape immunogenically hot tumors, which are associated with better response to therapy and improved prognosis (4). In the antigen presentation pathway, the interactions between peptide/major histocompatibility complex (MHC) and T cell receptors (TCR) are critical in triggering adaptive immunity (5). The MHC is highly polymorphic, enabling the presentation of a wide variety of antigens on the cell surface (6). In particular, the classical MHC I molecules (HLA-A, HLA-B, HLA-C) present endogenous antigens to CD8 T cells, and MHC II molecules (HLA-DR, HLA-DQ, HLA-DM) present exogenous antigens to CD4 helper T cells (7,8). The less polymorphic non-classical MHC I molecules (HLA-E, HLA-G) can act as inhibitory ligands to NK cells and contribute to immune tolerance (9). The non-classical MHC II molecules (HLA-DM, HLA-DO) act as chaperones and regulate antigen processing and loading (10). In the TME, both the generation of tumor-specific antigens (neoantigens) and the expression of antigen-presenting MHC molecules are important for effective immunity. A large pool of neoantigens derived from tumor mutations help trigger T cell response, shaping the formation of hot tumors (11,12). Deep learning algorithms have been built to predict high-affinity MHC-neoantigen binding, which contributes to the development of neoantigen-based cancer vaccines (13). On the other hand, immunohistochemistry (IHC) results have shown that upregulation of Human Leukocyte Antigen (HLA) class I expression in early-stage tumors may lead to CD8 T cell-mediated anti-tumor immunity (14). Inducement of HLA I expression in tumors by pro-inflammatory cytokines or by the inhibition of DNA methyltransferase can also result in strong cytotoxic CD8 T cell response (15,16).

Although these studies have demonstrated the potential role of HLA genes and neoantigen presentation in triggering immune-mediated tumor rejection, pan-cancer HLA class I and class II gene-mediated immunogenicity and tumor escape has not been systematically studied. Here, we aimed to uncover how HLA class I and II differential regulation shapes various immune microenvironments in 33 TCGA tumor types. While HLA supertypes showed an unbiased distribution across cancer types, HLA-B allele groups were associated with more variable survival outcomes. HLA expression was positively correlated with immune characteristics, and this correlation was even stronger than tumor mutational load, a metric widely used to evaluate immunogenicity (17). In addition, HLA allelic diversity in patients with high HLA expression was associated with improved survival. While upregulated HLA I expression was observed in most tumors with high immune infiltration, upregulated HLA II expression was uniquely seen in a subset of immunogenically hot tumors, and was correlated with helper T cell-mediated immunity and improved survival. In contrast, tumors with upregulation of HLA I but not HLA II genes showed DNA hypermethylation near HLA I genes and HLA LOH under strong immunity, suggesting that they had undergone immunoediting and exploited escape mechanisms. We further proposed machine learning models trained on HLA class I and class II gene expression data to predict the featured immune microenvironments with high accuracy. Our results emphasize the crucial role of HLA gene upregulation in triggering effective immunity and demonstrate the potential clinical applications using HLA expression data to evaluate tumor immunogenicity and predict prognosis.

## Materials and Methods

### Study Cohort and Data Acquisition

The samples analyzed in this study include primary tumors of 33 TCGA tumor types. Gene-level RNA-Seq expression data, methylation array files from Illumina Infinium Human Methylation 450k (HM450), and whole-exome sequencing bam files of tumors and matched tumor-adjacent normal samples were obtained from GDC Data Portal (https://portal.gdc.cancer.gov/).

The GTEx project includes samples from healthy individuals for over 50 tissues. In this study, we chose GTEx samples from 16 tissues where TCGA tumor-adjacent normal samples were also available. Gene-level RNA-seq expression data and phenotype metadata were acquired from UCSC Xena Treehouse (https://xenabrowser.net/datapages/?cohort=TCGA%20TARGET%20GTEx).

For analysis of HLA differential expression between tumor and matched normal tissues, 24 tumor types with available RNA-seq expression data of normal tissues were included. For analysis of tumor-infiltrating immune cells and progression-free survival, LAML was excluded for its property of not forming solid tumors and having no available PFS data (18). For analysis of HLA allelic divergence, LAML, ESCA, OV were excluded due to insufficient coverage at HLA class I loci for typing. For analysis of immune subtypes, LAML, THYM, and DLBC were excluded because their subtypes were not computed by Thorssen et al. (19).

### HLA Gene Expression

FPKM reads of RNA-seq expression data were obtained. A tissue-normalized FPKM value at each gene *X* was calculated as

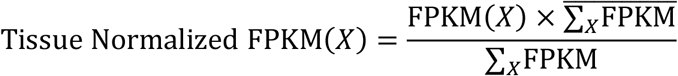

Where ∑_*X*_FPKM represents the sum of FPKM reads across all genes in a given sample, and 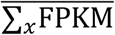 represents the tissue-specific sample average of the sum of FPKM reads across all genes. The sample average was each measured in 33 TCGA tumor types and the normal tissues, and was used to normalize the gene expression data of the sample belonging to that tissue type.

Individual gene expression was represented as the normalized log 2 value of tissue-normalized FPKM reads. Expression of HLA-DP, HLA-DQ, HLA-DR, HLA-DM, HLA-DO were represented as

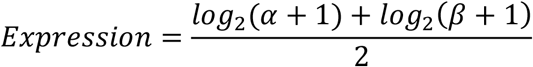

where *α* and *β* represents the transcript levels of *α* and *β* subunits, respectively. Overall tumor and normal HLA class I gene expression were measured as the log-transformed geometric mean of *B2M, HLA-A, HLA-B*, and *HLA-C* expression. Tumor and normal HLA class II gene expression were measured as the log-transformed geometric mean of *HLA-DRA, HLA-DRB1, HLA-DQA1, HLA-DQB1, HLA-DPA1*, and *HLA-DPB1* expression.

HLA class I gene expression fold change was represented as 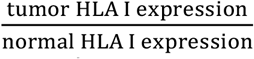. HLA class II gene expression fold change was represented as 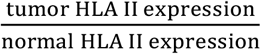.

### HLA Typing

The patient-specific 4-digit HLA types were acquired from Thorssen et al. (19), who used OptiType software (20) taking tumor RNA-seq fastq files as input. The HLA types of patients not typed previously were inferred using HLA-HD (21) taking whole-exome sequencing BAM files as input.

### HLA Gene Methylation in Tumor and Normal Samples

DNA methylation of HLA genes was measured as 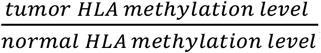, where HLA methylation level was represented as the mean β-scores across all CpGs near HLA I gene loci from the HM450 array. For each sample, HLA genes were hypermethylated if 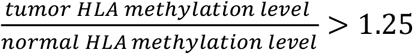, or hypomethylated if 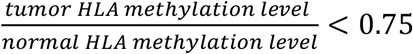.

### Immune Subtypes, Immune Cellular Fraction Estimates, Tumor Mutational Burden, and Cytolytic Activity

The immune subtypes, immune cellular fraction estimates, tumor mutational burden, and neoantigen load were obtained from Thorssen et al. (19). In brief, five immune signature modules were identified from 160 immune expression signatures, including macrophages/monocytes (22), overall lymphocyte infiltration (23), TGF-β response (24), IFN-*γ* response (25), and wound healing (26). These modules were further used to cluster 6 resulting immune subtypes C1-C6. To our knowledge, HLA/B2M genes were not used among the signatures for immune subtype clustering.

Immune cellular fraction estimates of 22 immune cell types were inferred with CIBERSORT (27) using TCGA RNA-Seq data.

For tumor mutational burden, protein-coding somatic mutation including insertions/deletions, missense mutations, nonsense mutations, frameshift mutations, nonstop mutations, splice site mutations, and transcription start site mutations were called. Nonsilent mutation rate per Mb was calculated.

For neoantigen prediction, neoepitopes were identified from single nucleotide variants (SNVs) and insertion-deletion mutations (Indels) using NetMHCpan v3.0 (28) with default settings. HLA calls and mutant peptides were used as inputs, and mutant peptides were identified as potential neoantigens if the predicted binding affinity (IC_50_) to autologous MHC < 500 nM and gene expression > 1.6 transcripts-per-million (TPM).

Cytolytic activity in each tumor sample was represented as cytolytic score (CYT) as previously proposed (29). The expression levels of *GZMA* (ENSG00000145649) and *PRF1* (ENSG00000180644) were used. The following formula was applied to represent CYT:

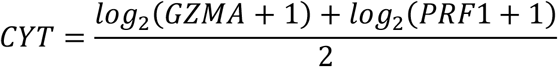

### Functional Divergence of HLA Alleles and Patient HED

HED was calculated as previously described (30). In brief, protein sequence of exons 2 and 3, namely the peptide-binding domain, of each HLA allele was extracted. Protein sequences and exon annotations were obtained from the ImMunoGeneTics/HLA (IMGT/HLA) database (https://www.ebi.ac.uk/ipd/imgt/hla/). Pairwise alignment between the protein sequences of two alleles in each HLA-I locus was performed using MUltiple Sequence Comparison by Log-Expectation (MUSCLE) (31). The Grantham distance was used to measure functional divergence between two alleles taking into account the physicochemical properties of amino acids. It was calculated as the sum of amino acid differences (including the biochemical composition, polarity, and volume of each amino acid) in the pairwise alignment along the protein sequence of peptide-binding domains following the formula by R. Grantham (32):

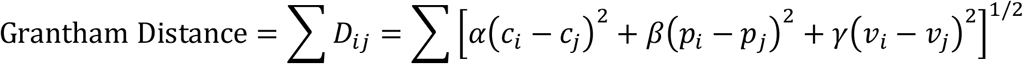

where *D* is the Grantham distance between the aligned sequences, and *i* and *j* are the amino acids at a homologous position. *c, p*, and *v* represent biochemical composition, polarity, and volume of the amino acids, respectively. *α, β*, and *γ* are constants as originally proposed. The Grantham distance was normalized by the length of the sequence alignment. Patient HED was calculated as the mean of Grantham distances at *HLA-A, HLA-B*, and *HLA-C*.

### Principal Component Analysis

The expression of classical HLA class I genes (*HLA-A, HLA-B, HLA-C*), *B2M*, non-classical HLA class I genes (*HLA-E, HLA-G*), classical HLA class II genes (*HLA-DRA, HLA-DRB1, HLA-DQA1, HLA-DQB1, HLA-DPA1, HLA-DPB1*), and non-classical HLA class II genes (*HLA-DMA, HLA-DMB, HLA-DOA, HLA-DOB*) was averaged within individual tumor type, respectively. The features were centered and projected onto principal components (PCs).

### Classification

To classify tumors with high cytolytic activity, defined as the top quantile of cytolytic score, one-hot encoding was applied on TCGA tumor type, and log 2 values of classical and non-classical HLA class I (*HLA-A, HLA-B, HLA-C, HLA-E, HLA-G*), *B2M*, and class II gene (*HLA-DRA, HLA-DRB1, HLA-DQA1, HLA-DQB1, HLA-DPA1, HLA-DPB1, HLA-DMA, HLA-DMB, HLA-DOA, HLA-DOB*) expression in tumors were used to train a random forest classifier with n_estimators=200 and criterion=‘gini’.

To predict the immune subtypes of all tumors, one-hot encoding was applied on TCGA tumor types; the log 2 values of *B2M* and HLA gene expression in tumors were used to train a three-layer neural network model with hyperbolic tangent activation function, Adam optimizer, Sparse-Categorical-Cross-Entropy loss function, and accuracy metric. The input layer contains 128 neurons; the hidden layer contains 96 neurons; the output layer contains 6 neurons with SoftMax activation function.

For all classifications, the dataset was divided into training sets (60%) and testing sets (40%) randomly. Area under the curve (AUC) was calculated to evaluate the performance of the classifiers. For multiclass classification, a micro-average AUC was computed to account for class imbalance and evaluate the overall performance.

### HLA Loss of Heterozygosity (LOH) Analysis

LOHHLA (Loss of Heterozygosity in Human Leukocyte Antigen) (33) was used to assess HLA LOH in BRCA, LUAD, and SKCM patients. The program requires a tumor and germline BAM, patient-specific HLA typing, HLA allele sequence FASTA file, and purity and ploidy estimates. Purity and ploidy estimates were acquired from GDC (https://portal.gdc.cancer.gov/). The estimated allele 1 and allele 2 copy number for each patient (HLA_type1copyNum_withBAFBin, HLA_type2copyNum_withBAFBin, respectively), as well as a p-value (PVal_unique), were extracted from the LOHHLA output file.

A patient is identified as fully heterozygous if the individual has distinct alleles at all loci of *HLA-A, HLA-B, HLA-C*. The classification of HLA LOH in a tumor was the same as originally proposed by (33). In specific, a tumor sample was identified as harboring HLA LOH if an allele’s copy number was less than 0.5 with p value less than 0.05. The number of allelic loss for each patient was calculated as the total number of alleles being classified as LOH.

### Statistics and Survival Analysis

Statistical analyses were performed in R. Wilcoxon Rank Sum Test and Kruskal-Wallis Test were used for comparison of a variable of interest in two groups and multiple groups, respectively. Two-sided Fisher’s Exact Test was used to determine the significance of any difference in proportions between two classifications. Spearman’s rank correlation was used to determine the correlation of two variables.

The values of progression-free interval (PFI) and status were used as obtained from (18). To examine the survival effect of a continuous variable x, samples were split equally into high-x and low-x patients and analyzed using log-rank test and Kaplan-Meier estimator. In HED score analysis, the variable x represents HED score and HLA class I gene expression. In HLA class II expression fold change analysis across individual tumor types, the variable x represents HLA class II fold change in tumor samples with defined immune subtypes. In HLA LOH analysis, samples were split into groups with high HLA I expression or low HLA I expression, and the survival efficacy of HLA LOH within each group was assessed.

To compare the performance of cytolytic activity, tumor mutational burden, neoantigen load, HLA I fold change and HLA II fold change on survival prediction, to compare the four groups defined based on HLA I expression and HED score, and to compare the four groups defined based on HLA I and II expression fold change, Cox proportional-hazards modeling was used.

## Lead Contact

Further information and requests for resources and reagents should be directed to and will be fulfilled by the lead contact, Rachel Karchin.

## Material Availability

This study did not generate new unique reagents.

## Data and Code Availability

- This paper analyzes existing, publicly available data. Gene expression data and downstream analysis data are available within the article and its supplementary data files.
- All original code has been deposited at https://github.com/KarchinLab/HLA-Mediated-Tumor-Immunogenicity-Manuscript and is publicly available as of the date of publication.
- Any additional information required to reanalyze the data reported in this paper is available from the lead contact upon request.

## Results

### HLA genes Are Differentially Expressed in Most TCGA Tumor Types, with the Expression of Class I Genes Higher than Class II

We first assessed individual HLA class I and class II gene expression, as well as Beta-2-microglobulin (*B2M*) in association with MHC I heavy chains, in all 33 TCGA tumor types. While most HLA class I genes are expressed ubiquitously, HLA class II expression is restricted to professional antigen-presenting cells (APCs) (34). Overall, *B2M* and most HLA class I genes demonstrated higher expression than class II genes (Figure 1A). Consistent with the fact that HLA-G and HLA-DO molecules have more tissue-restricted distribution (35,36), we observed the lowest expression in *HLA-G* and HLA-DO (*HLA-DOA, HLA-DOB*) among class I and class II genes, respectively.

**Figure 1.**
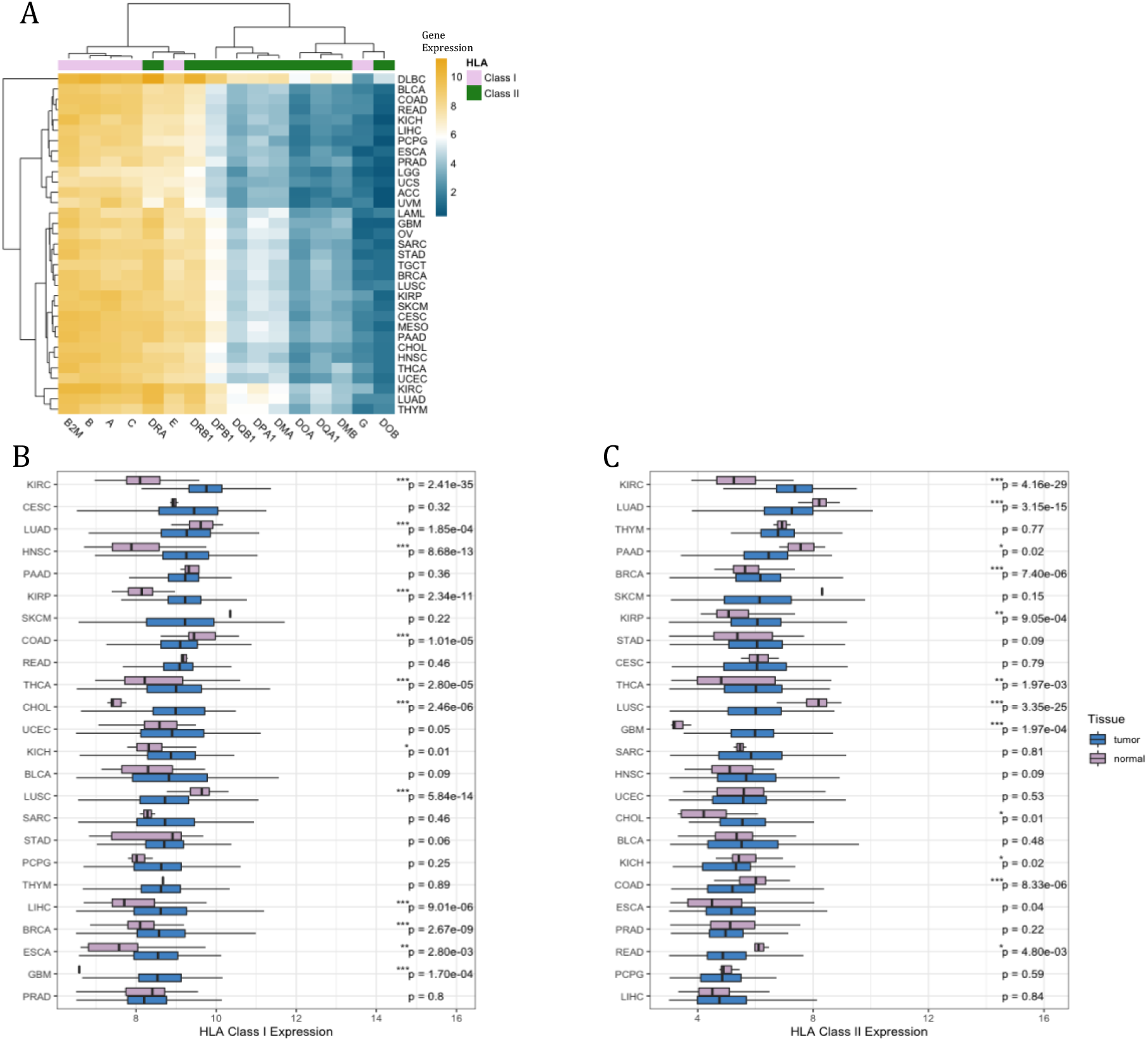
HLA Expression Is Differentially Expressed across Tumor Types. (A) Expression of classical and non-classical HLA class I and class II genes in tumor samples (n=11080) across 33 TCGA tumor types. Expression of gene *X* is represented as the mean gene expression of all samples within the tumor type. (B) HLA class I expression, and (C) HLA class II expression in tumor-normal pairs. Boxes in the box plot represent interquartile ranges and horizontal lines represent 5^th^-95^th^ percentile ranges. p values between the pairwise tumor-normal samples are calculated by Wilcoxon rank-sum test and are unadjusted. Asterisks denote significant FDR-adjusted (Benjamini & Hochberg method) p values (*p<0.05; **p<0.01; ***p<0.001).

Next, we evaluated HLA I and II differential expression between tumor and normal samples. 10 tumor types had significantly upregulated HLA class I expression and 3 had downregulated class I expression, while 6 tumor types had upregulated class II expression and 6 had downregulated class II expression (Figure 1B-C). Among them, KIRC showed the most upregulation of HLA class I genes and class II genes, while LUSC had the most downregulation of HLA genes. The variability in HLA gene differential expression is likely due to tissue of origin and reflects a difference in tumor-specific mechanisms to regulate HLA gene expression.

For reference, we also compared HLA class I and II expression between TCGA tumors and GTEx normal samples of the same tissue type. Many TCGA tumors downregulated HLA I expression, with LUSC and LUAD showing the most downregulation (Figure S1A). Yet, most TCGA tumors had significantly higher HLA II expression than GTEx normal samples (Figure S1B). Since GTEx normal samples are from healthy donors, it is possible that TCGA tumors and tumor-adjacent normal tissues are in an immunosuppressive microenvironment, resulting in systematic downregulation of HLA I expression. In contrast, HLA II genes are primarily expressed by APCs, which aggregate in lymph nodes and spleens in healthy individuals. Upon infection or chronic inflammation, APCs are recruited to the site of inflammation and form tertiary lymphoid organs (TLOs) in the periphery for local antigen presentation and T cell activation (37). Hence, GTEx healthy tissues may contain fewer APCs than inflamed tumors and show lowered overall HLA II expression.

### HLA Class I and Class II Gene Expression Correlate with Tumor Immune Characteristics and Predict Cytolytic Activity

We then evaluated whether HLA expression was associated with immune characteristics in tumors, including immune infiltration, proinflammatory gene signatures, and immune checkpoint signatures. Both HLA class I and class II expression demonstrated a strong correlation with anti-tumor immune infiltration (Figure 2A). They were also positively correlated with the expression of pro-inflammatory genes, as well as immune checkpoints. Intriguingly, HLA gene expression displayed a stronger correlation with these immune characteristics than TMB or neoantigen load, which are commonly used to predict immunogenicity and therapy response (13,17). In addition, while TMB and neoantigen load showed weak correlation with cytolytic activity (CYT), a metric previously proposed by Rooney et al. (29) as the log average of Granzyme A (*GZMA*) and Perforin-1 (*PRF1*), the positive correlation between HLA expression and CYT was stronger (Spearman correlation, TMB: R=0.21; neoantigens: R=0.24; HLA I: R=0.62; HLA II: R=0.74) (Figure S2A).

**Figure 2.**
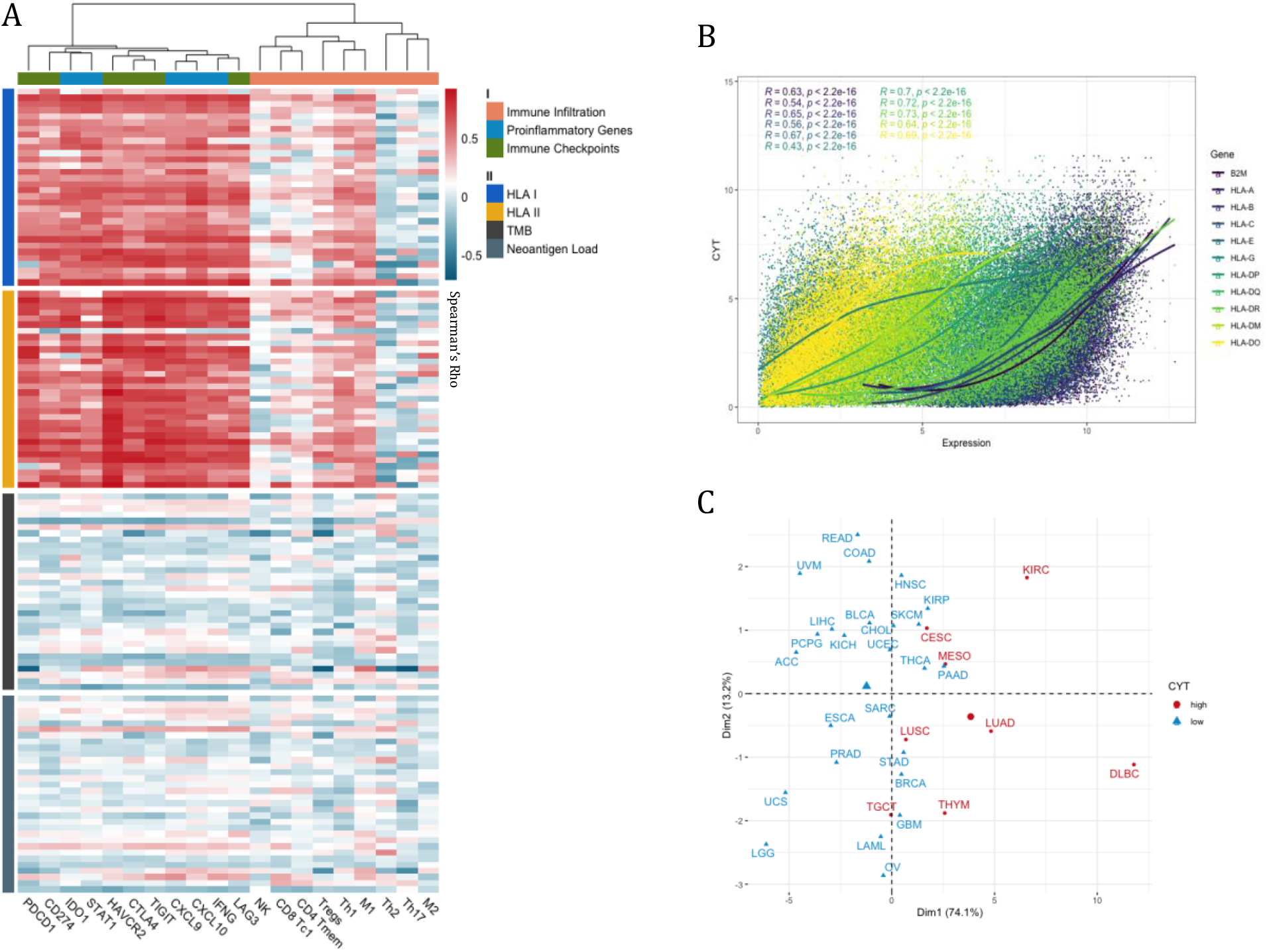
HLA Class I and Class II Gene Expression Are Associated with Immune Activity. (A) Spearman correlations between HLA class I, class II expression, TMB, neoantigen load, and immune characteristics. Each row within a horizontal block represents a distinct tumor type. CD8 Tc1: CD8 cytotoxic T cells; CD4 Tmem: activated CD4 memory T cells; Th1: T helper 1 cells; Th2: T helper 2 cells; Th17: T helper 17 cells; Tregs: regulatory T cells; NK: natural killer cells; M1: Macrophage 1; M2: Macrophage 2. (B) Spearman correlations between the expression of individual HLA genes and cytolytic activity (CYT). p values are calculated from Spearman’s rank correlation and are unadjusted. (C) Principal Component Analysis (PCA) on 33 tumor types by their expression of classical and non-classical HLA class I and class II genes. Principal Component 1 (Dim1) and Principal Component 2 (Dim2) are shown. High cytolytic activity (CYT high) is defined as the top 25% of mean CYT.

Among the classical and non-classical HLA I genes, *HLA-E* demonstrated the highest correlation with CYT (R=0.67), while *HLA-G* showed the lowest (R=0.43) (Figure 2B). Among HLA II genes, HLA-DR showed the highest correlation with CYT (R=0.73), while HLA-DM had the lowest (R=0.64). An unsupervised principal-component analysis (PCA) of all HLA I and HLA II expression data showed that tumors with high cytolytic activity (top quantile) clustered together (Figure 2C). In addition, we trained a random forest classifier using as features classical and non-classical HLA I and HLA II gene expression, and were able to classify samples showing strong cytolytic activity with high accuracy (AUC=0.93), suggesting that the expression data has potential for clinical prediction of immunogenically hot tumors (Figure S2B). Consistent with the strength of their correlation with CYT, *HLA-E, HLA-DQA1* and *HLA-DRA* demonstrated the highest relative feature importance as reported by the classifier, while *HLA-G* had the lowest (Figure S2C). Taken together, HLA class I and II expression were strongly associated with tumor immunogenicity in our study and could potentially outperform TMB as a clinical metric to predict immune activity.

### HLA Allelic Diversity in Tumors with High HLA Expression Correlates with Improved Survival

MHC-I molecules are highly variable with divergent biophysiochemical properties in the peptide-binding grooves to present antigens. We next assessed the distribution of HLA class I supertypes (38,39) among TCGA tumors. Overall, the supertypes showed an unbiased distribution across different tumors (Figure 3A). A24 was the least predominant HLA-A supertypes, while B8, B27, B58 and B62 were least predominant among HLA-B supertypes. HLA I expression did not show significant variation across the supertypes, suggesting that the specific HLA types of an individual were independent of HLA gene expression (Figure S3A-C). In addition, immune infiltration, proinflammatory gene expression, and immune checkpoint expression were comparable between different HLA supertypes (Figure S3D-E). Among HLA I genes, HLA-B allele groups were associated with more variable patient survival, with B*15 (Cox proportional-hazards, HR=1.79, p=0.021) and B*50 correlated with significantly worse patient survival (HR=21.15, p=0.003) (Figure S4A-C). Since HLA-B was the most diverse of all three HLA I loci (40), the results emphasized that HLA-B alleles might contribute more to varied patient prognosis by facilitating the presentation of divergent epitopes.

**Figure 3.**
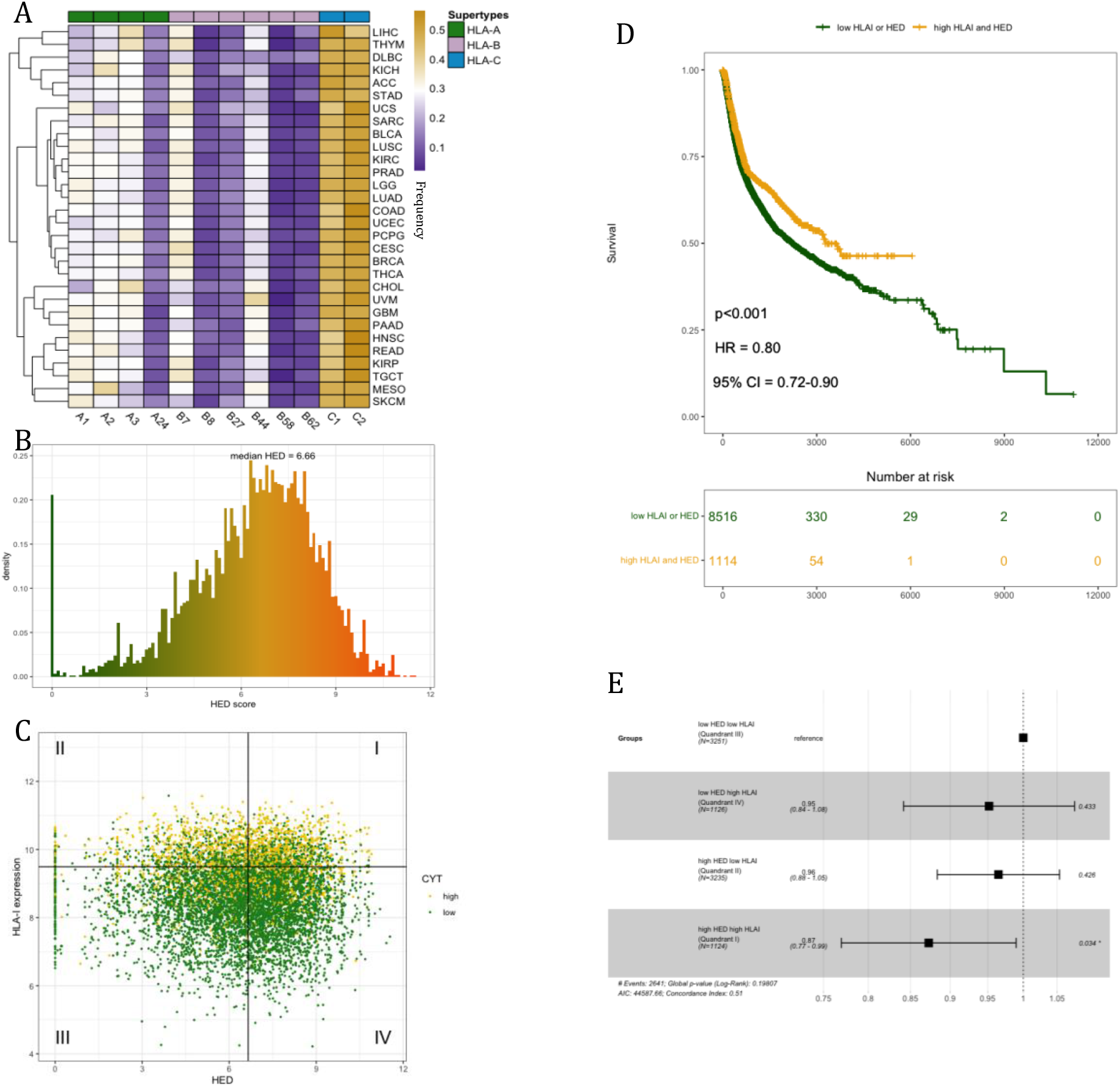
HLA-I Evolutionary Divergence (HED) in Tumors with High HLA Expression Is Associated with Improved Survival. (A) Frequencies of HLA class I supertypes among tumor types. Color scales represents the frequency of occurrence for each supertype in the corresponding tumor types. (B) HED score distribution from 30 tumor types (n=8859). (C) HED score vs. HLA class I expression. High cytolytic activity (CYT high) is defined as the top 25% of all tumor samples. The vertical line represents the median HED score, and the horizontal line represents the 75^th^ percentile HLA class I expression. Four quadrants are defined accordingly. (D) Progression-free survival (PFS) by high HLA I expression and HED (Quadrant I) and low HLA I expression or HED (Quadrant II, III, IV). p-value, HR, and 95% confidence interval are calculated by the Log-rank test. (E) The hazard ratio (HR) of the effects of HED and HLA I expression on PFS from Cox Proportional-hazards model, with low HED low HLAI as reference. Squares represent the HR and horizontal bars represent 95% confidence intervals. HED: HLA-I evolutionary divergence score; HLAI: HLA class I expression.

Since a higher diversity of a patient’s HLA alleles could potentially allow a larger pool of neoantigens to be presented and hence increase the likelihood of the tumor being rejected by immunity (30), we asked if HLA allelic diversity is correlated with tumor immunogenicity. The biophysiochemical diversity of each patient’s HLA class I alleles was estimated with the HLA-I evolutionary divergence (HED) score (41). A patient homozygous at all three HLA I loci has a HED score of 0. The greater the functional diversity of HLA I alleles, the higher the patient’s HED score (Figure S3F). The median HED of all patients was 6.66, with a low being 0 and a high being 11.54 (Figure 3B). However, we did not observe any correlations between HLA I allelic diversity and infiltration levels of immune cells, pro-inflammatory genes, or immune checkpoints in the TME (Figure S3G).

We reasoned that high allelic diversity might not contribute to the presentation of a large neoantigen pool if HLA expression is low. Hence, we proposed that high allelic divergence of HLA I alleles in tumors with high HLA I expression may more efficiently present neoantigens and lead to tumor rejection. To test this hypothesis, we divided all patients into four quadrants by their HED score and HLA I expression (Figure 3C). Patients within Quadrant I with high HLA I expression and high HED scores demonstrated significantly better survival (log-rank test, HR=0.80, p<0.001) (Figure 3D). Moreover, among the four quadrants, high HLA I (Quadrant IV) or high HED (Quadrant II) alone did not show a significant correlation with survival (Figure 3E). We concluded that high HLA I allelic diversity in tumors with high HLA I expression was associated with better survival.

### HLA Class I and Class II Genes Show Differential Expression Patterns across Various Immune Microenvironments with Different Survival Outcomes

Next, we asked if HLA genes were differentially regulated within various immune microenvironments in tumors. To this end, we evaluated HLA class I and II differential gene expression in tumors compared to their matched normal samples across six immune subtypes (C1-C6) previously clustered from 160 immune expression signatures, to our knowledge excluding HLA genes: C1 (wound healing), C2 (IFN-*γ* dominant), C3 (inflammatory), C4 (lymphocyte depleted), C5 (immunologically quiet), and C6 (TGF-β dominant) (19). C2 and C3 are characterized by anti-tumor Type I immune response. C5 is distinguished by the lowest level of immune infiltrates, while C4 and C6 have the worst prognosis with a macrophage-dominated TME (19). We found that HLA class I expression was upregulated in C2 and C3 (Wilcoxon rank-sum, p<2e-16), and was downregulated the most in C5 (p=0.028) (Figure 4A, 4C). This is consistent with the characterization of C2 and C3 having the highest immune infiltration and C5 having the lowest. HLA class II expression was significantly downregulated in C1 (p=5.1e-11), C4 (p=0.0038), and C5 (p=0.032) (Figure 4B-C). Surprisingly, though both C2 and C3 are immunogenically hot, HLA II was uniquely upregulated in C3 (p=8.8e-13), suggesting that the upregulation of HLA I alone and upregulation of both HLA classes may shape the immune microenvironment differently.

**Figure 4.**
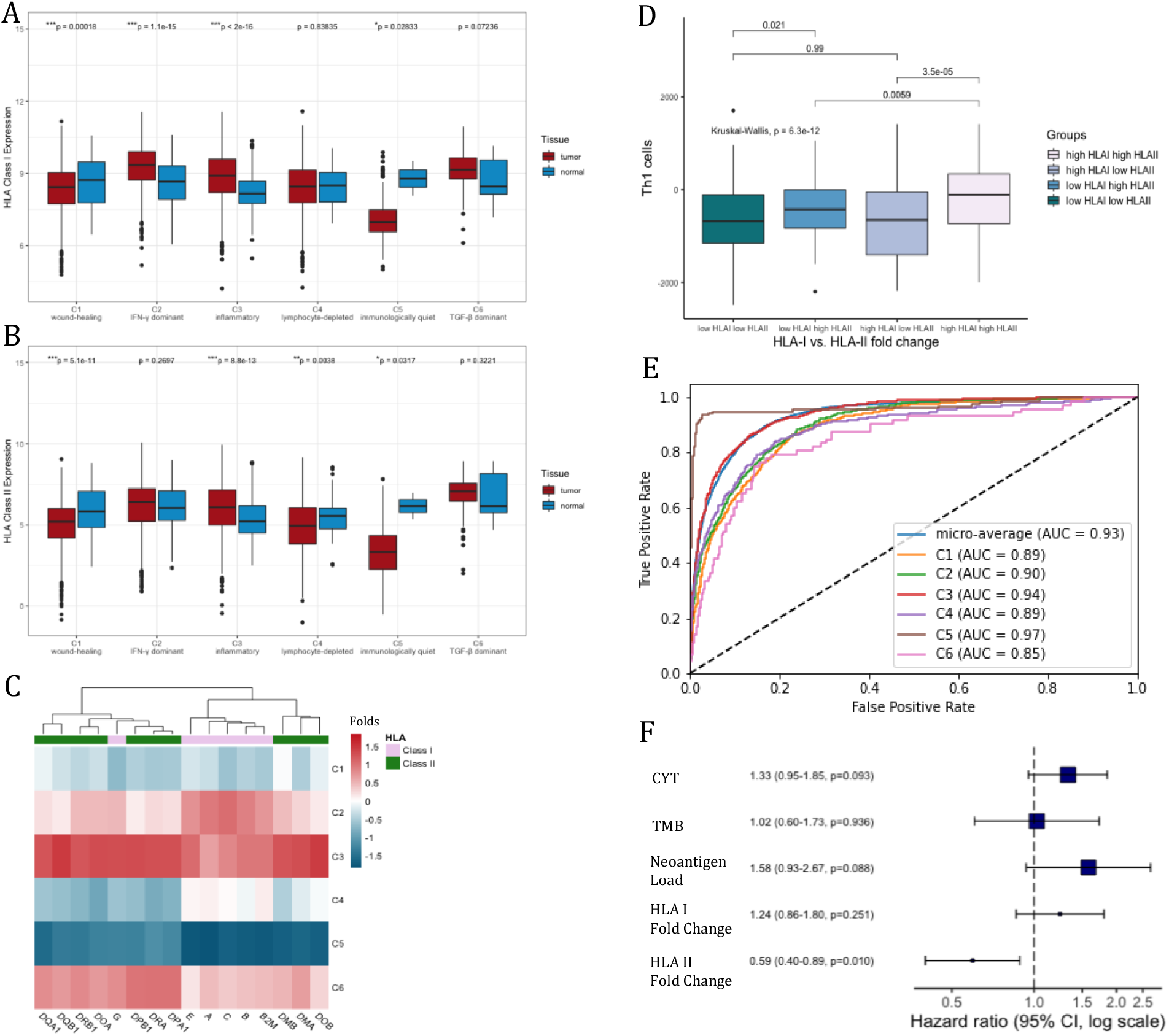
HLA Expression Predicts the Immune Subtypes and Is Associated with Improved Survival. (A) HLA class I expression, and (B) HLA class II expression between tumors and matched normal samples across six immune subtypes (n=9126), C1-C6. Boxes in the box plot represent interquartile ranges. P values between pairwise tumor-normal samples are calculated by Wilcoxon rank-sum test. Asterisks denote significant FDR-adjusted (Benjamini & Hochberg method) p values (*p<0.05; **p<0.01; ***p<0.001). (C) Expression fold change of HLA class I and class II genes across six immune subtypes. The heatmap is scaled by columns. (D) Helper 1 T cells (Th1) signatures associated with HLA class I and class II expression fold change. High HLA I and high HLA II are defined as the top 50% of all tumor samples, respectively. Pairwise p values are calculated by Wilcoxon rank-sum test. p-value across all 4 groups is calculated by the Kruskal-Wallis test. (E) Receiver Operating Characteristic (ROC) curve of a neural network model. Micro-average area under the curve (AUC) as well as AUC for individual immune subtype predictions are presented. (F) The hazard ratio (HR) of CYT, TMB, neoantigen load, HLA I fold change, and HLA II fold change on patient PFS from Cox Proportional-hazards model. Squares denote the HR and horizontal bars represent 95% confidence intervals.

Since HLA class I molecules interact with CD8 T cells, while class II molecules interact with CD4 helper T cells, we next evaluated if HLA class I and class II gene upregulation were correlated with helper vs. cytotoxic T cell-mediated adaptive immunity. Patients with high HLA-II fold change demonstrated significantly elevated Th1 cell fractions than those with low HLA-II fold change (Figure 4D). In comparison, high HLA I fold change only weakly correlated with increased Th1 cell fractions. On the other hand, high HLA-I and HLA-II fold change both correlated with increased cytolytic activity and CD8 T cell infiltration, yet they were more significantly elevated in groups with high HLA-II fold change (Figure S5A-B). While both the upregulation of HLA class I and class II genes were associated with cytotoxic T cell-mediated immunity, our results suggest a strong correlation between HLA class II upregulation and helper T cell-mediated immunity, which might lead to more effective T cell priming and stronger cytolytic activity in the TME.

We considered if HLA gene expression data could be used to predict the immune subtype of a given tumor sample. An artificial neural network model was trained on the transcriptomic data of HLA class I and class II genes to predict the six immune subtypes. The model was capable to predict the immune subtype of tumors with high accuracy (micro-average AUC=0.93) (Figure 4E). Among the six immune subtypes, tumors in C3 and C5 were classified with the highest accuracy (C3: AUC=0.94, C5: AUC=0.97), while tumors in C6 were classified with the lowest accuracy (C6: AUC=0.85). Overall, the results demonstrated that HLA class I and class II gene expression were highly variable and had the potential to predict different types of immune microenvironments in tumors.

We then asked whether HLA expression fold change could predict patient survival. While the commonly used metrics to predict prognosis, including cytolytic activity (Cox proportional hazards, HR=1.33, p=0.093), TMB (HR=1.02, p=0.94), and neoantigen load (HR=1.58, p=0.088), performed comparably, HLA II fold change demonstrated significantly better prognosis (HR=0.59, p=0.010) (Figure 4F). Also, among individual tumor types, high HLA-II fold change within LUAD and BRCA was associated with significantly better survival (Log-rank test, LUAD: HR=0.31, p=0.011; BRCA: HR=0.43, p=0.048) (Figure S5C). High HLA-II expression fold change in other tumor types (KIRC, THCA, COAD, UCEC, ESCA, HNSC, LIHC, BLCA) also correlated with better survival, though the correlation was not statistically significant. We concluded that upregulation of HLA class II genes was correlated with effective immune response and rejection of tumors, potentially contributing to better prognosis.

### DNA Methylation near HLA Class I Genes Favored under Strong Cytolytic Activity Can Be Alleviated by Class II Gene Upregulation

DNA methylation near HLA genes has previously been shown to downregulate gene expression and dampen the immune response as a potential HLA-mediated tumor escape strategy (15). We first evaluated the effect of differentially methylated HLA class I genes on immunity. Overall, there existed a significant inverse correlation between HLA I expression and methylation levels (Spearman correlation, R=-0.61, p<2.2e-16). Tumors with high cytolytic activity were more hypomethylated near HLA I genes, whereas those with low cytolytic activity were hypermethylated (Figure 5A). Moreover, we observed varied HLA I methylation levels across the immune subtypes (Kruskal-Wallis, p<2.2e-16) (Figure 5B). Consistent with low HLA I gene expression observed in C1 (wound healing) and C4 (lymphocyte depleted), HLA I methylation was elevated in C1 and C4. Surprisingly, tumors in C2 (IFN-*γ* dominant) demonstrated significantly higher HLA I methylation compared to tumors in C3 (inflammatory) (Wilcoxon rank-sum, p=9.5e-6), though their HLA I expression fold change was comparable (Figure S6A). This suggests that while tumors in both C2 and C3 upregulated HLA I expression, C2 tumors were at the same time heavily methylated near HLA I genes.

**Figure 5.**
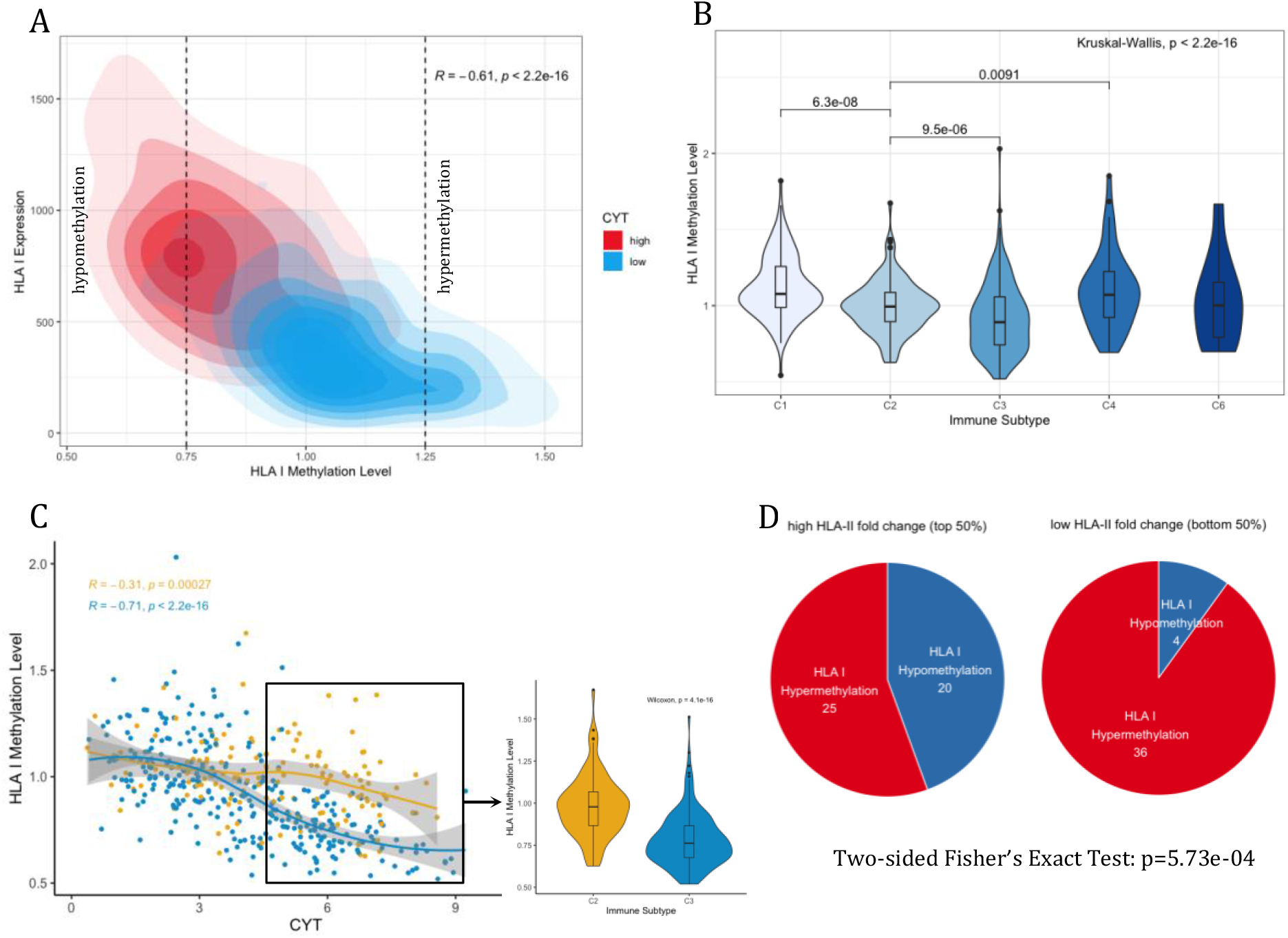
HLA Class I Methylation Is Favored in A Subset of Hot Tumors and Dampens Immune Activity. (A) A kernel density plot of HLA class I methylation levels and expression across groups (n=657) with high (top 25%) or low cytolytic activity. Correlation coefficient and p values are calculated from Spearman’s rank correlation. CYT: cytolytic activity; Hypomethylation: methylation < 0.75; Hypermethylation: methylation > 1.25. (B) HLA class I methylation levels across five immune subtypes (excluding C5 with no available data). p values between pairwise immune subtypes are calculated by Wilcoxon rank-sum test and are unadjusted. p-value across all immune subtypes is calculated by the Kruskal-Wallis test. (C) Left panel: The Spearman correlations between cytolytic activity (CYT) and HLA class I methylation in C2 and C3 tumors. Right panel: HLA class I methylation in tumors with high cytolytic activity (top 50%) across C2 and C3. p-value is calculated by Wilcoxon rank-sum test and is unadjusted. (D) Association between HLA II fold change and aberrant methylation states near HLA I genes in all samples. Numbers on pie charts indicate the number of tumor samples showing hypermethylated or hypomethylated HLA I genes. p-value is calculated from two-sided Fisher’s Exact Test.

We specifically focused on tumors in C2 and C3, which were both defined by Type I immune response (19). C2 tumors with high cytolytic activity showed significantly heavier HLA I methylation than C3 tumors (Wilcoxon rank-sum, p=4.1e-16), whereas tumors with low cytolytic activity showed comparable HLA I methylation (Figure 5C). This suggests that a subset of tumors with Type I immune response have increased HLA I methylation to evade recognition under strong immune activity. Since HLA class II genes were significantly upregulated in C3 (Figure S6A), we asked if higher HLA II fold change would alleviate HLA I methylation in tumors. Tumors with high HLA-II fold change were significantly more hypomethylated than those with low HLA-II fold change (Two-sided Fisher’s Exact Test: p=5.73e-04) (Figure 5D). Our results support that HLA class II gene upregulation may be a mechanism to reduce tumor escape and trigger effective immunity in the TME.

### HLA Loss of Heterozygosity (LOH) Is Associated with Worse Survival but Can Be Counter Balanced by High HLA Expression

Lastly, we considered HLA LOH, another potential HLA-mediated tumor escape strategy, and evaluated how it might correlate with HLA expression and patient survival. We focused on BRCA, LUAD and SKCM, three common cancer types with potential for immunotherapy. Overall, HLA LOH correlated with worse survival (Log-rank test, HR=1.28, p=0.043) (Figure S7A). Among BRCA and SKCM patients, HLA LOH did not show a significant correlation with survival (BRCA: HR=0.66, p=0.11; SKCM: HR=0.36, P=0.2), whereas in LUAD patients HLA LOH was correlated with worse survival (HR=1.42, p=0.020) (Figure S7B-D). The results suggest that HLA LOH may associate with varied tumor characteristics and trigger different immune mechanisms across cancer types.

Next, we proposed that tumors with HLA LOH under strong immune selection were more favored to evade recognition. Supporting our hypothesis, tumors with high cytolytic activity harbored more HLA LOH than those with low cytolytic activity (Two-sided Fisher’s exact Test, p=9.52e-8) (Figure 6A). Across the five immune subtypes, tumors in C2 (IFN-*γ* dominant) demonstrated the highest percentage of HLA LOH, while tumors in C3 (inflammatory) had the lowest (BRCA C2: 23%, C3: 12%; LUAD C2: 41%, C3: 21%; SKCM C2: 21%, C3: 7%) (Figure 6B). Consistent with our observations of HLA I methylation, though the tumors in both C2 and C3 were characterized with Type I immune response, a larger fraction of C2 tumors harbored HLA LOH, suggesting potential immunoediting and tumor escape.

**Figure 6.**
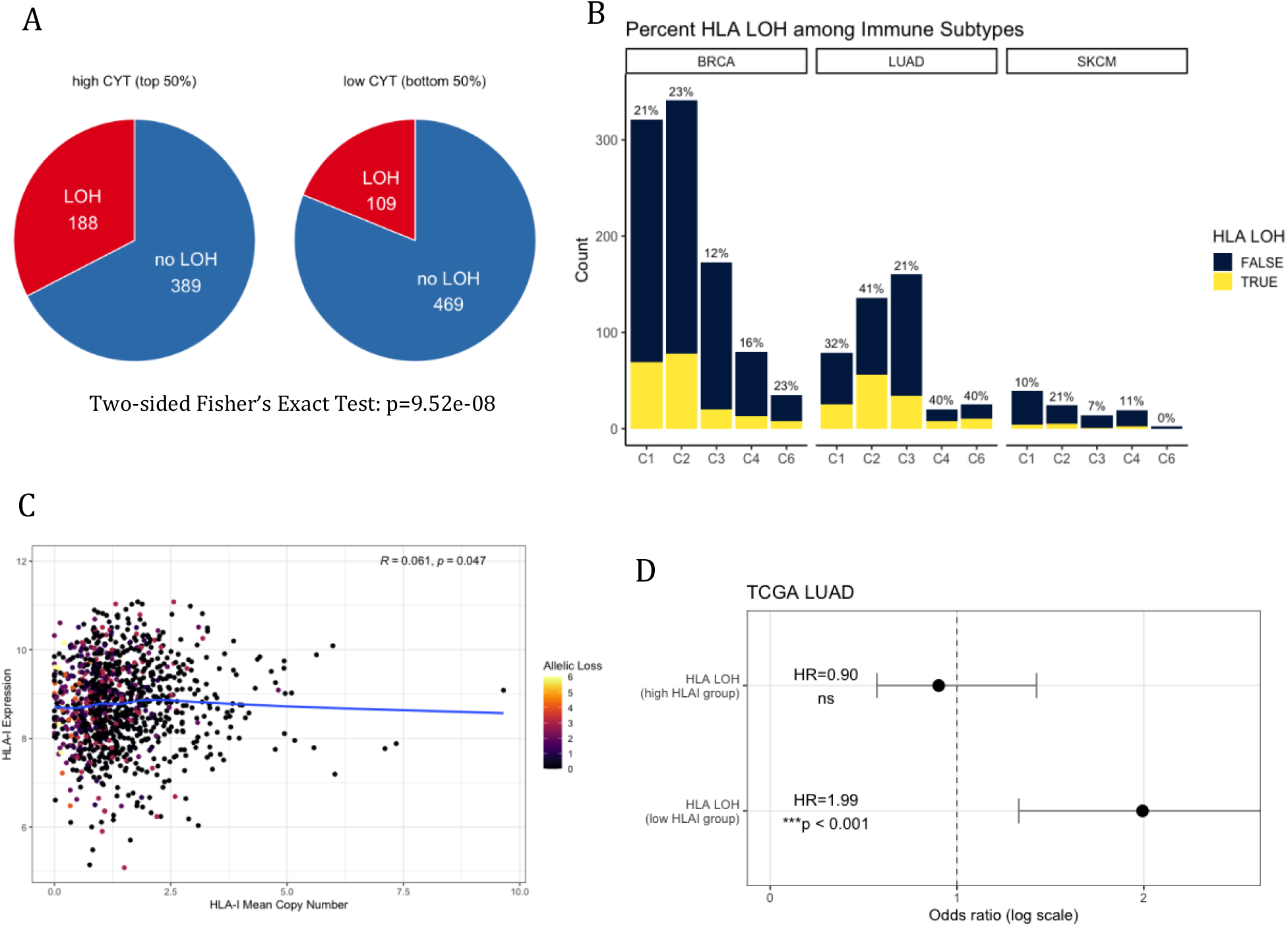
HLA LOH Is Exploited by Hot Tumors with Worse Patients’ Survival. (A) Association between cytolytic activity and HLA LOH. Numbers on pie charts indicate the number of tumors harboring HLA LOH. Patients with heterozygosity at all three HLA-I loci are included (n=1155). p-value is calculated from two-sided Fisher’s Exact Test. (B) Tumor samples harboring HLA LOH across five immune subtypes (excluding C5 with no available data) in BRCA (n=932), LUAD (n=419) and SKCM (n=98), respectively. Percentage on top of each bar represents the percent of tumor samples with HLA LOH within the corresponding immune subtype. (C) Spearman correlation between the mean copy number of HLA class I genes and HLA class I expression. The allelic loss represents the number of lost HLA I alleles. (D) The hazard ratio (HR) of the effect of HLA LOH on LUAD patient PFS among high HLA-I expressing tumors and low HLA-I expressing tumors from log-rank test. Circles represent the HR and horizontal bars represent 95% confidence intervals. High HLAI group: high HLA class I expression (top 50%); low HLAI group: low HLA class I expression.

Surprisingly, we did not find any correlation between the mean HLA I copy number and gene expression (Spearman correlation, R=0.061, p=0.047) (Figure 6C, S7E-G). We also did not find association between HLA I expression or expression fold change and LOH (Figure S7H-I), as previously proposed by Garrido et al. (5). This suggests that HLA I expression at the RNA level is uncorrelated with HLA I allelic loss at the DNA level. HLA LOH most likely does not downregulate total HLA expression as a means of facilitating tumor escape.

Based on the notion that HLA LOH may lead to a loss of the most immunogenic neoantigens being presented on the cell surface and result in worse survival (33), we explored if high HLA gene expression may compensate for reduced neoantigen presentation, leading to worse prognosis in LUAD. Interestingly, worse patient survival was observed in low HLA-I expressing tumors with HLA LOH (Log-rank test, HR=1.99, p<0.001) but not in high HLA-I expressing tumors with HLA LOH (HR=0.90, ns) (Figure 6D). We concluded that the correlation between HLA LOH and worse survival may be counterbalanced by higher HLA expression.

## Discussion

Based on the idea that antigen-presenting HLA class I and II molecules are fundamental for triggering anti-tumor immunity, we aimed to address how HLA expression shapes various immune microenvironments and tumor escape in a pan-cancer analysis of tumor types from TCGA. Our analysis suggests that elevated HLA expression is likely to drive the formation of immunogenic tumors. Our results also demonstrate the predictive power of HLA gene expression data on the immune microenvironment and survival.

While mutational load is frequently used to predict immunogenicity and survival, HLA I and HLA II expression showed much stronger correlation with immune characteristics. We reasoned that neoantigens generated in tumors with high TMB but low HLA expression could not be effectively presented to trigger immune activities. Hence, high HLA expression is necessary for effective antigen presentation, and it likely plays a greater role in inducing immune activity than tumor mutations. In recent years, neoantigen-based personalized cancer vaccines and adoptive cell therapy (ACT) are among the advanced immunotherapies to trigger neoantigen-specific T cell activity (42). Current *in silico* algorithms predict neoantigens based solely on a patient’s HLA types and tumor mutations (43,44). Here, we suggest that HLA expression is a crucial component that should be incorporated in neoantigen prediction algorithms.

We further demonstrated that the expression levels of classical and non-classical HLA genes could be used to train a probabilistic classifier to predict tumors with high cytolytic activity in association with tumor immunogenicity (29). Interestingly, although the non-classical HLA-E molecule can act as an inhibitor against NK cells (45,46), it showed the strongest correlation with cytolytic activity among class I genes and was also the most important feature in the Random Forest classification. HLA-E is known to play a role in both innate and adaptive immunity. Whereas it inhibits NK-mediated lysis by interacting with the CD94/NKG2A complex on NK cells, it can present a specific pool of peptides to activate HLA-E restricted CD8 T cells independent of the classical HLA-restricted T cells (46). Future efforts can be made to explore the specific roles of HLA-E molecules in anti-tumor adaptive immunity. Our results suggest that inducing HLA expression in tumors systematically may trigger prolonged anti-tumor immunity and enhance the efficacy of immunotherapies.

We also considered HLA allelic diversity, which potentially correlates with the diversity of neoantigens being presented (41). We found that high HLA allelic diversity in tumors expressing high levels of HLA genes correlated with improved survival. This suggests that high HLA allelic diversity may contribute more to tumor elimination through the presentation of a diverse neoantigen pool when HLA expression is high. Yet, the role of HLA allelic diversity alone in triggering immune activities is limited. One possible explanation is that high HLA gene expression despite low allelic diversity may still trigger immunity by presenting sufficient amounts of neoantigens even if the presented neoantigens are of low diversity. In contrast, high HLA allelic diversity, though having the potential to present a more diverse group of neoantigens, cannot compensate for low HLA expression in cells with reduced neoantigen presentation.

HLA genes were differentially regulated across six immune subtypes. Tumors with Type I immunity showed higher HLA expression fold change than non-immunogenic tumors. While HLA I molecules expressed on tumors present endogenous neoantigens to CD8 cytotoxic T cells, most HLA II molecules expressed on professional APCs present exogenous neoepitopes to CD4 helper T cells (47). CD4 T cell priming is essential in activating CD8 cytotoxic T cells and forming prolonged memory (48,49). As expected, high HLA II fold change was correlated with helper T cell-mediated immunity, as well as elevated immune cytotoxicity, leading to better prognosis. We suggest that HLA II fold change may be a better metric than tumor mutational load for predicting prognosis. Taken together, the results support the crucial role of HLA class II upregulation in T cell priming and generation of strong cytotoxicity for tumor elimination. We further demonstrated that HLA class I and class II gene expression could predict the immune subtypes of tumors with high accuracy. The neural network model uses 15 HLA gene expression signatures as features and could potentially help oncologists diagnose the tumor immune microenvironment and propose personalized therapies for better prognosis. Future work could include training classifiers on HLA expression data of immunotherapy-treated patients to predict response to therapy.

Strong immune selective pressure can lead to tumor immunoediting. Here, we considered DNA methylation near HLA I genes and HLA LOH as potential HLA-mediated immune evasion strategies. A subset of immunogenic tumors with high cytolytic activity showed more hypermethylated HLA genes, potentially as a means to reduce neoantigen presentation and evade recogonition. High HLA-II fold change was associated with more hypomethylated HLA I genes, whereas low fold change was associated with hypermethylated genes. Hence, we suggest that, in addition to its role in triggering effective immunity, HLA II upregulation reduces immunoediting through HLA gene methylation. In contrast, tumors with high HLA I fold change alone may experience strong T cell cytotoxicity in the absence of helper T cell activation, leading to immunoediting and eventually tumor escape.

Finally, we assessed the role of HLA LOH in anti-tumor immunity. HLA LOH correlated with varied survival outcomes across cancers. While previous studies propose that HLA I allelic loss results in tumors evading immune recognition potentially through loss of allele-specific expression (5,33), we did not observe any correlation between HLA LOH and HLA downregulation. This suggests that HLA LOH likely results in tumor evasion not through downregulating the overall HLA expression, but by other mechanisms such as reducing the presentation of the most immunogenic neoantigens on the cell surface due to allele-specific HLA loss. Furthermore, we observed that higher expression of HLA genes could counterbalance the negative impact of HLA LOH on survival. We propose that high HLA expression may allow higher numbers of neoantigens to be presented and effectively compensate for immune evasion through reduced presentation of the most immunogenic neoantigens, leading to tumor rejection and improved survival. Since the overall HLA expression was not correlated with HLA LOH, this finding suggests a new way of rescuing the anti-tumor immune recognition in patients with HLA LOH. Although HLA LOH, unlike HLA gene methylation, is an irreversible hard lesion at the gene level, therapies that enhance the expression of HLA genes at the RNA and protein levels could be exploited. Possible future work includes inducing HLA gene expression with pro-inflammatory cytokines like IFN-*γ* in tumors with HLA LOH to evaluate if elevated gene expression rescues immune recognition and tumor elimination *in vivo*.

Our study has several limitations. First, the HLA expression metric we used was transcript-based and did not take into account post-translational modification, thus might not accurately capture the final quantity of HLA molecules expressed on the cell surface. In addition, though next-generation sequencing has made practical HLA typing from short-read data with high resolution, high polymorphism and homology of HLA genes still pose significant challenge for typing (21). We also did not consider specific neoantigen pools in tumors that could bind to the HLA molecules and be presented to T cells. Nonetheless, our analysis of TCGA samples has emphasized the important role of HLA differential regulation in facilitating the formation of prolonged immune response in tumors with improved survival. We also demonstrate the power of HLA expression data in predicting various immune microenvironments. Our framework should be useful for cancer immunotherapy studies and for understanding immunoediting mechanisms.

## Acknowledgments

The results published here are in part based upon data generated by the TCGA Research Network: https://www.cancer.gov/tcga.

The Genotype-Tissue Expression (GTEx) Project was supported by the Common Fund of the Office of the Director of the National Institutes of Health, and by NCI, NHGRI, NHLBI, NIDA, NIMH, and NINDS. The data used for the analyses described in this manuscript were obtained from: the GTEx Portal and/or dbGaP accession number phs000424.vN.pN.

## Author Contributions

XG conceived the study and performed the experiments, XG and RK wrote the paper.

**Figure S1.**
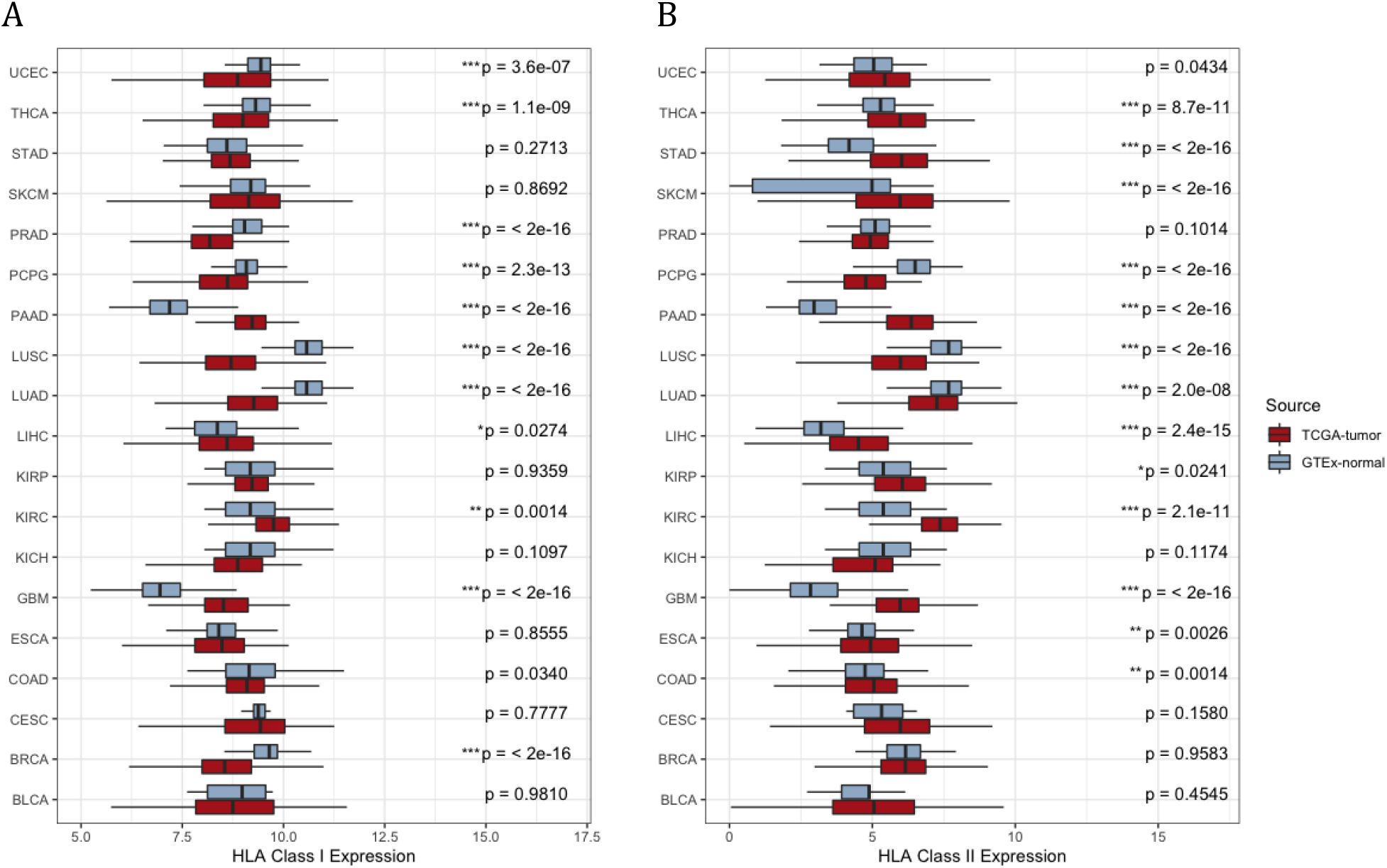
HLA Gene Expression Compared between Tumors and Tissue-Matched Healthy Donors, related to Figure 1. (A) HLA class I expression, and (B) HLA class II expression, between TCGA tumor samples and GTEx normal samples across tissues. p values between the pairwise tumor-normal samples are calculated by Wilcoxon rank-sum test and are unadjusted. Asterisks denote significant FDR-adjusted (Benjamini & Hochberg method) p values (*p<0.05; **p<0.01; ***p<0.001).

**Figure S2.**
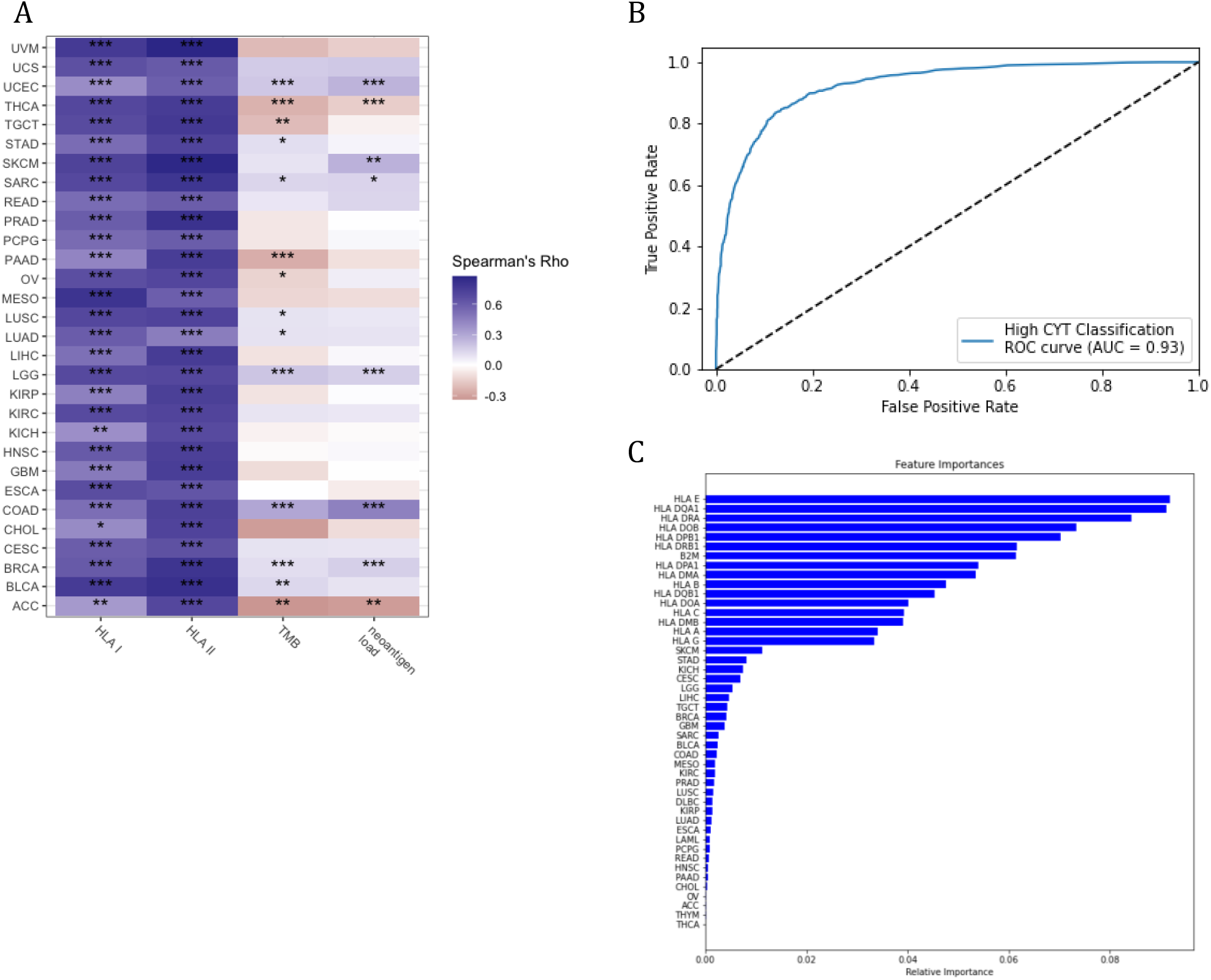
HLA Class I and Class II Expression Data Predict Cytolytic Activity with High Accuracy, related to Figure 2. (A) Spearman correlations between HLA class I, class II expression, TMB, neoantigen load, and cytolytic activity across tumor types. Asterisks denote significant p values (*p<0.05; **p<0.01; ***p<0.001). (B) Receiver operating Characteristic (ROC) curve of a random forest classifier. The random forest classifier is trained on log-transformed expression levels for classical and non-classical HLA class I and class II genes and predicts tumors with high cytolytic activity (high CYT). AUC: area under the curve. (C) The relative importance of classical and non-classical HLA gene expression in the random forest classifier. All features used to train the classifier are listed on the y-axis.

**Figure S3.**
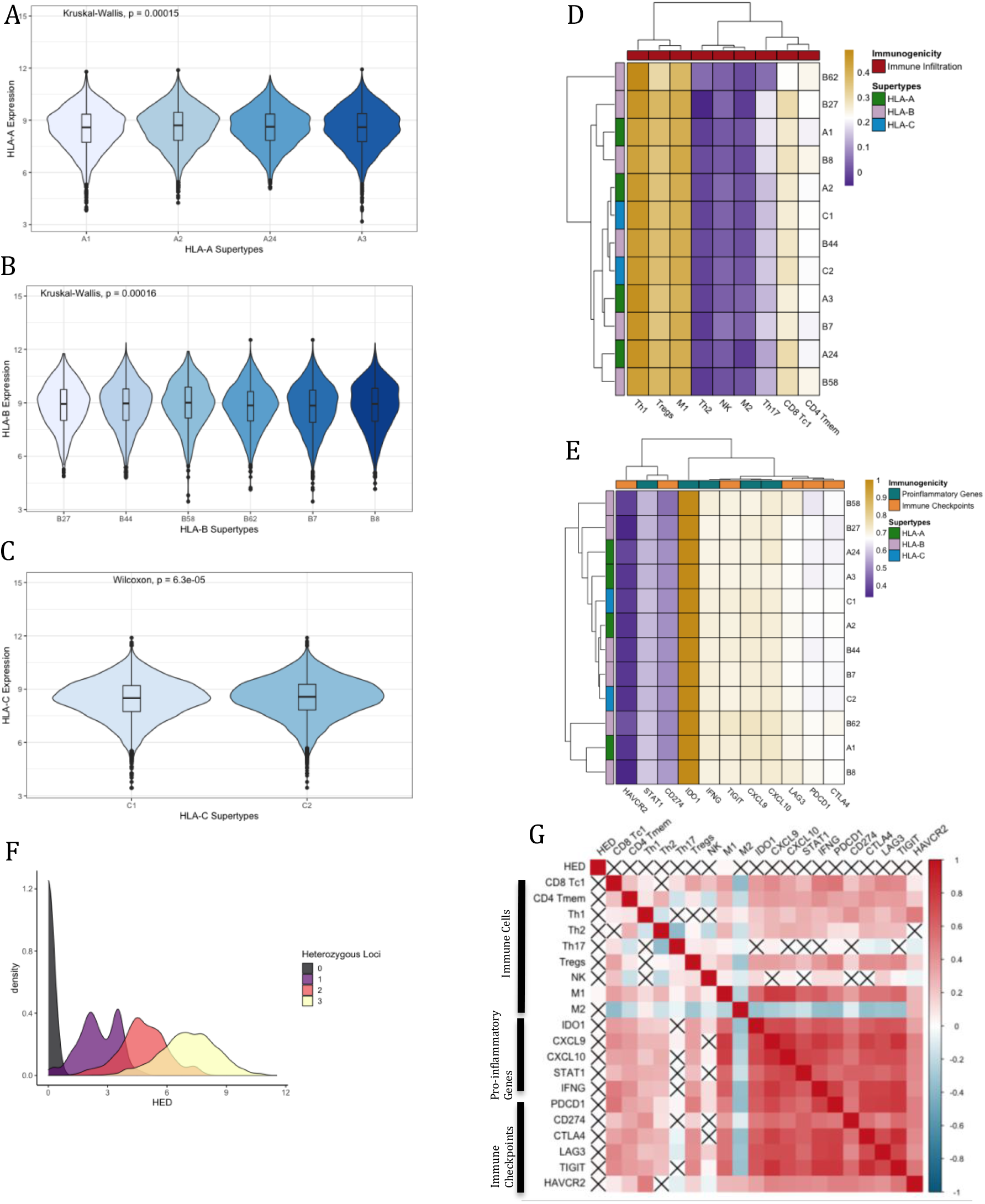
HLA supertypes Are Uncorrelated with Immune Characteristics, related to Figure 3. (A) The association between HLA-A supertypes and HLA-A expression, (B) HLA-B supertypes and HLA-B expression, and (C) HLA-C supertypes and HLA-C expression. p value is calculated by Kruskal-Wallis test. (D) Immune infiltration, represented as the relative fraction of each immune cell type, across HLA class I supertypes. (E) The expression of proinflammatory genes and immune checkpoints across HLA class I supertypes. (F) HED score distribution across groups of patients with different number of heterozygous HLA I loci. (G) Spearman’s rank correlation between HED score and immune characteristics. Unadjusted two-tailed p values are reported below the diagonal and adjusted p values (Benjamini & Hochberg method) for multiple testing are reported above the diagonal. Correlations not significant (p > 0.01) are marked with an X in each corresponding cell.

**Figure S4.**
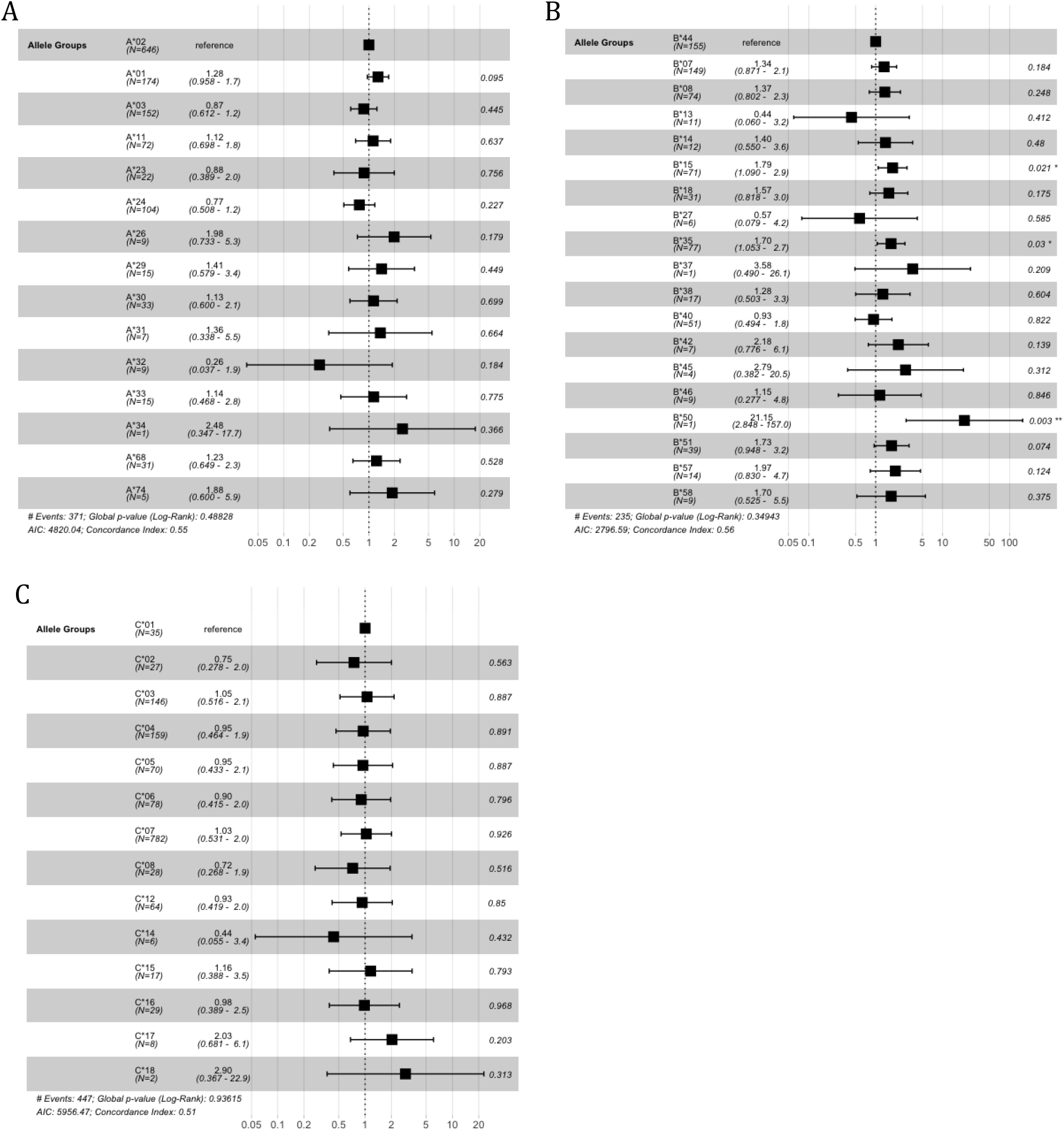
HLA Class I Allele Groups Associate with Variable Patient Survival, related to Figure 3. (A) The hazard ratio of the effects of HLA-A allele groups, (B) HLA-B allele groups, and (C) HLA-C allele groups, on progression-free survival (PFS). A*02, B*44, C*01 were the most common allele groups among all patients and were used as the reference in each Cox-proportional hazards model, respectively. Only patients with homozygosity at the corresponding locus (HLA-A: n=1295, HLA-B: n=738, HLA-C: n=1451) are included. A*25, B*39, B*48, B*49, B*52, B*53, B*54, B*55, B*81 are excluded in the analysis for having less than 10 cases and all patients discontinued or ceased at the time of assessment. HR: hazard ratio; CI: confidence interval.

**Figure S5.**
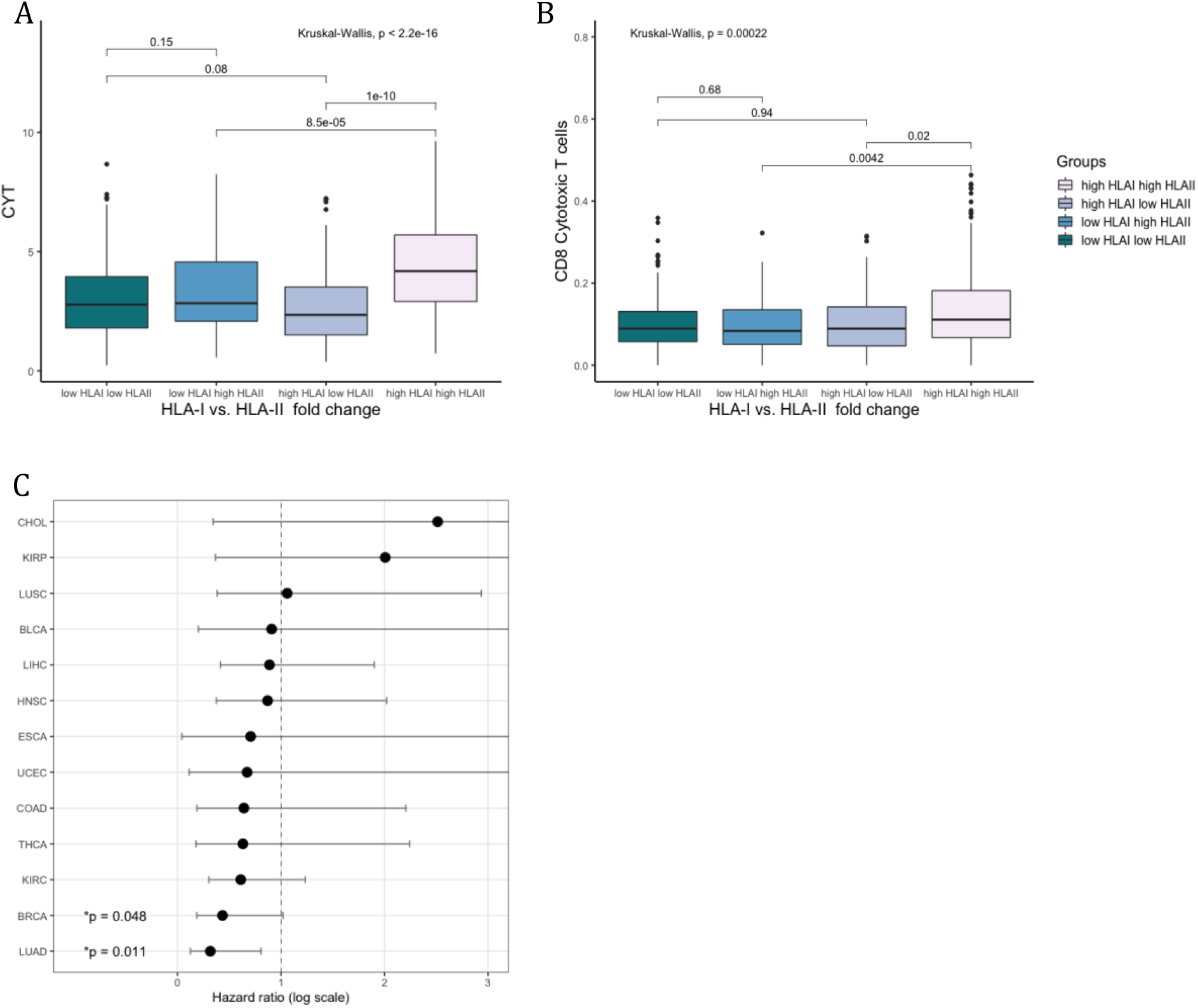
HLA Class II Upregulation Correlates with Strong Immunity and Improved Survival, related to Figure 4. (A) Cytolytic activity (CYT) across 4 groups based on their HLA class I and class II expression fold change. High HLA I and high HLA II are defined as the top 50% of all tumor samples, respectively. Pairwise p values are calculated by Wilcoxon rank-sum test. p-value across all 4 groups is calculated by the Kruskal-Wallis test. (B) Same as (A), but for infiltration levels of CD8 cytotoxic T cells. (C) The hazard ratio (HR) of the effect of high HLA class II expression fold change (top 50%) on PFS across 13 TCGA tumor types with more than 5 cases of tumor/normal pairs available. Circles represent the HR and horizontal bars represent 95% confidence intervals. HR and p values are calculated from the log-rank test.

**Figure S6.**
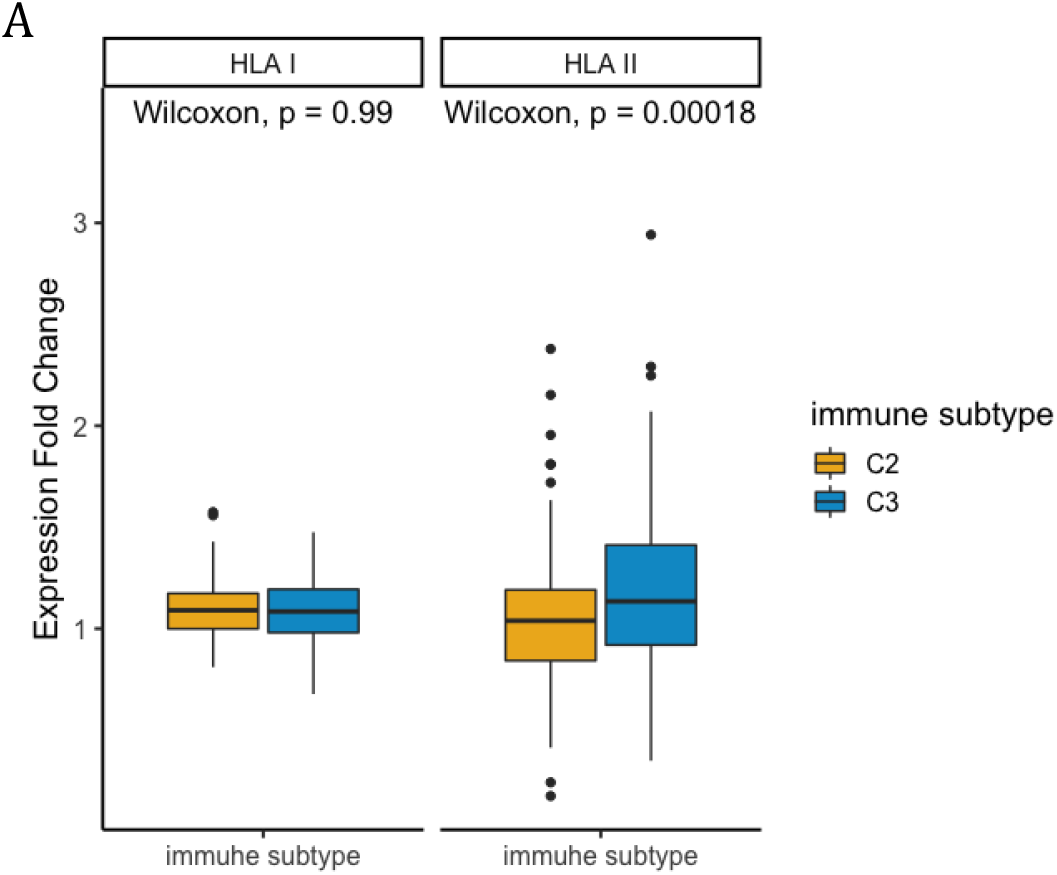
The Immune Subtypes of Type I Immune Response Showed Differentially Expressed HLA Class I and Class II Genes, related to Figure 5. (A) HLA class I and class II expression fold change are compared between C2 (n=2591) and C3 (n=2397) tumors. p values are calculated by Wilcoxon rank-sum test and are unadjusted.

**Figure S7.**
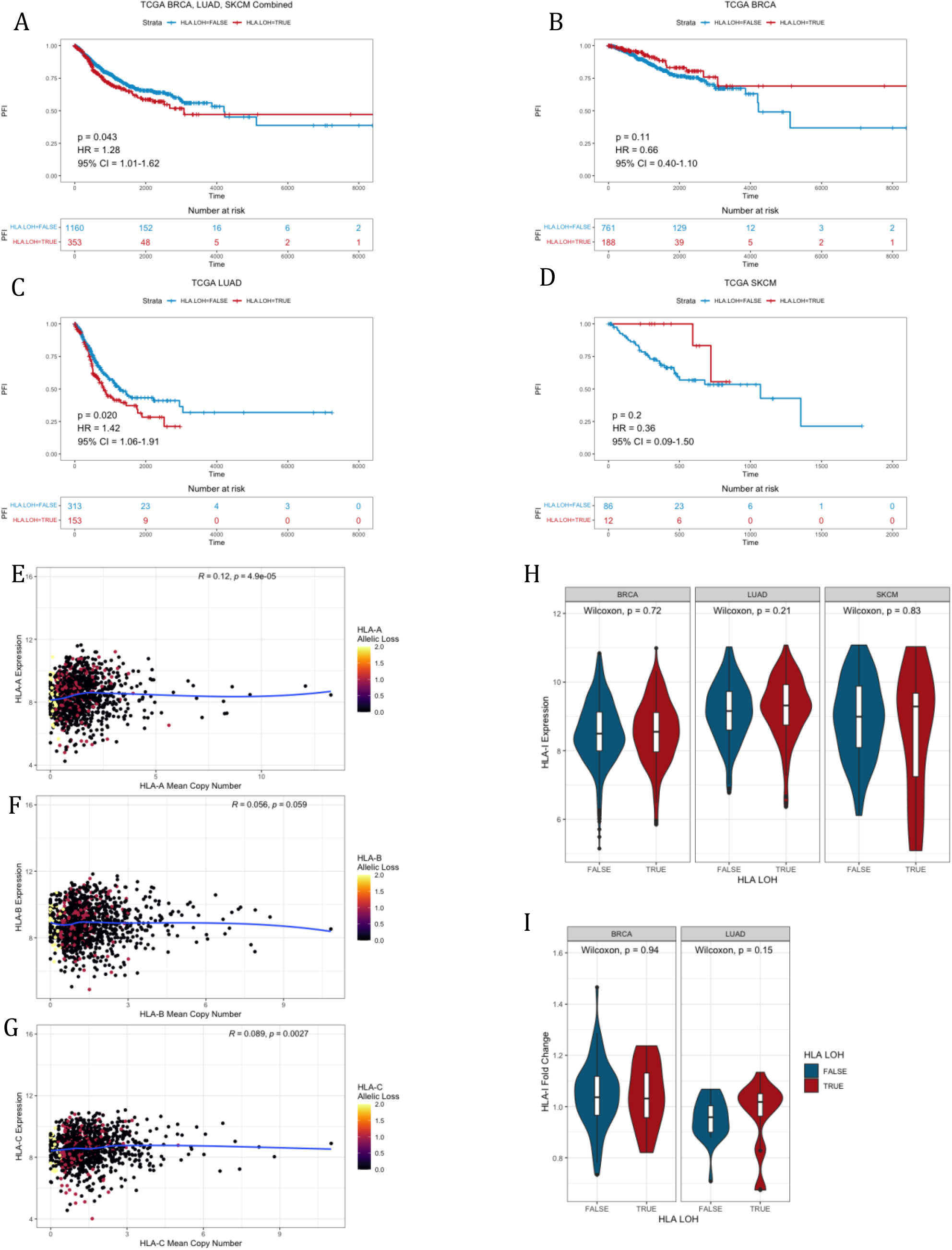
HLA LOH Contributes to Worse Survival but Does Not Downregulate HLA Gene Expression, related to Figure 6. (A) Progression-free survival (PFS) by HLA LOH among all BRCA, LUAD, SKCM patients (n=1513), (B) among BRCA patients (n=949), (C) among LUAD patients (n=466), and (D) among SKCM patients (n=98). (E) Spearman correlation between the mean copy number of *HLA-A*, (F) *HLA-B*, (G) *HLA-C*, and expression. The correlation coefficient and p-value are calculated from Spearman’s rank correlation. The allelic loss represents the number of lost HLA I alleles. (H) The association between HLA class I expression and HLA LOH. (I) The association between HLA class I expression fold change and HLA LOH. SKCM is excluded for having only one case with expression fold change. p-value is calculated by Wilcoxon rank-sum test and is unadjusted.

## Supplementary Data

**Supplementary File 1. TCGA Tumor/Normal Samples Availability and Clinical Data, related to Figure 1, 4, 6**.

**Supplementary File 2. HLA/B2M Signatures Expression in TCGA and GTEx Samples, related to Figure 1**.

**Supplementary File 3. Immune Signatures Expression and Cytolytic Activity, related to Figure 2**.

**Supplementary File 4. HLA-I Supertypes and HLA-I Evolutionary Divergence (HED) Scores, related to Figure 3**.

**Supplementary File 5. DNA Methylation Beta-Values of HLA Genes, related to Figure 5**.

**Supplementary File 6. HLA Loss of Heterozygosity (LOH), related to Figure 6**.

## Notes

### Competing Interest Statement

The authors have declared no competing interest.

